# Structural Pockets and Interacting RNA-Associated Ligands (SPIRAL): A DSSR-enabled Meta-Analysis of RNA-Small Molecule Recognition

**DOI:** 10.64898/2026.05.19.726393

**Authors:** Xiang-Jun Lu, Yaqiang Wang

## Abstract

Small molecules that target structured RNA hold therapeutic promise across a wide range of diseases, yet the structural principles governing RNA–ligand recognition remain poorly defined. We present SPIRAL (Structural Pockets and Interacting RNA-Associated Ligands), a curated database of 1,098 RNA–small molecule structures from the Protein Data Bank covering 1,137 ligand-binding events across six functional RNA categories. A customized pipeline built on DSSR (Dissecting the Spatial Structure of RNA) extracts structural interaction parameters from each complex, capturing stacking geometry, hydrogen-bond topology resolved by RNA moiety, groove engagement, and tertiary motif context. Unsupervised clustering of these fingerprints resolves six mechanistically distinct binding modes, the distribution of which is strongly governed by RNA functional class. To enable category-independent comparison of interaction quality across these diverse modes, we introduce the Composite Binding Quality Score (CBQS), a seven-metric framework that ranks riboswitches highest and regulatory RNA motifs lowest among the six categories. Across 275 affinity-characterized entries, C2′-endo sugar pucker count and total buried contact surface area emerge as the dominant predictors of binding affinity, converging with the structural features most underengaged by current regulatory RNA motif binders. SPIRAL provides a data-driven foundation for the rational design of next-generation RNA-targeted therapeutics.

## INTRODUCTION

Approved small-molecule drugs overwhelmingly target proteins, yet protein-coding sequences account for less than 2% of the human genome, leaving the vast majority of the transcriptome pharmacologically unexplored (1,2). Many disease-relevant proteins, including transcription factors and intrinsically disordered proteins, lack defined hydrophobic binding pockets and have resisted small-molecule intervention for decades, underscoring the need for alternative target classes (3). Structured RNAs carry out essential regulatory, catalytic, and scaffold functions across the transcriptome, and a growing body of structural data confirms that many disease-relevant RNAs harbor well-defined binding pockets accessible to small molecules, making the transcriptome a compelling and largely untapped therapeutic target space (4–6). The clinical approval of risdiplam for spinal muscular atrophy, which acts by directly engaging the SMN2 pre-mRNA splice site to promote exon 7 inclusion, established that potent and selective RNA-targeting small molecules are achievable, providing the first clinical proof of concept for RNA as a primary small-molecule therapeutic target (7,8).

Despite this progress, the structural principles governing RNA–small molecule recognition remain far less well understood than those of protein–ligand recognition, due to several interconnected challenges. First, RNA adopts a wider diversity of tertiary folds than is accessible to most computational pocket-detection algorithms, which were developed and calibrated for hydrophobic protein cavities and perform poorly on the polar, electrostatic, and shape-defined pockets that characterize RNA binding sites (9,10). Second, existing databases of RNA-binding ligands are organized around different aspects of RNA–ligand recognition, but none provides systematic tertiary structural context at the level of individual binding interactions (11–14). Third, experimental binding affinity data are scattered across the literature and have not been systematically integrated with structural interaction parameters, preventing the construction of quantitative structure–affinity models across RNA functional classes. Addressing all three challenges simultaneously requires an analysis framework that extracts tertiary structural context from atomic coordinates rather than secondary structure annotation alone, operates uniformly across the full structural diversity of RNA functional classes, and integrates curated experimental binding affinities with these parameters, a combination of capabilities that existing resources do not provide.

DSSR (Dissecting the Spatial Structure of RNA) is a comprehensive RNA structural analysis algorithm that automatically detects, annotates, and classifies complex tertiary structural features (15,16), and its atomic-level annotation of base pairs, multiplets (higher-order coplanar base associations), stacking interactions, backbone geometry, and tertiary motif context provides exactly the principled basis needed to address these three challenges. To enable systematic analysis of RNA–small molecule recognition, we extended DSSR with a new module (termed “ligand”) specifically designed to characterize RNA–ligand interactions, including quantification of stacking interactions by differential solvent-accessible surface area, hydrogen-bond classification by RNA moiety contacted, and groove-preference assignment based on the distribution of contacts across the major and minor groove edges. These newly developed interaction features operate on the same atomic coordinate framework as DSSR’s existing tertiary structure annotation, ensuring that each interaction parameter is reported in its full tertiary structural context. Building on this extended pipeline, we extracted a wide range of structural interaction fingerprints from each RNA–ligand complex. These parameters capture both the chemical character of each interaction, including stacking, hydrogen bonding, 2′-OH contacts, and backbone contacts, groove engagement, and the tertiary motif context in which that interaction occurs, including non-canonical pairs, base multiplets, hairpin loops, internal loops, junction loops, pseudoknots, coaxial stacks, and G-quadruplexes.

Leveraging this pipeline, here we present SPIRAL (Structural Pockets and Interacting RNA-Associated Ligands), a curated database of 1,098 RNA–small molecule structures drawn from the PDB, covering 1,137 ligand-binding events across six functional RNA categories: riboswitches, ribozymes, synthetic aptamers, G-quadruplexes, ribosomal RNA, and regulatory RNA motifs. We introduce the Composite Binding Quality Score (CBQS), a seven-metric framework built on these interaction fingerprints that quantifies RNA–ligand interaction quality independent of RNA functional category and ligand size, enabling mechanistically grounded comparison across the full structural diversity of the dataset. Using CBQS together with the DSSR-derived interaction fingerprints, we ask (i) whether RNA functional class governs binding mode at the structural level, (ii) whether interaction quality can be compared in a mechanism-aware manner across functionally distinct RNA classes, and (iii) which structural features of the RNA–ligand interface predict binding affinity. Our central finding is that C2′-endo sugar pucker count and total buried contact surface area are the dominant predictors of binding affinity, and that this C2′-endo enrichment is concentrated at junction loops, non-canonical base pairs, and base multiplet networks, precisely the tertiary structural features most underengaged by current disease-relevant RNA binders. This convergence establishes a unified structural criterion for improving both potency and selectivity in RNA-targeted drug discovery.

## MATERIALS AND METHODS

### SPIRAL dataset curation

All RNA–small molecule structures were retrieved from the PDB (https://www.rcsb.org, May 12, 2026) using the “ligand” module of DSSR v2.9.0, which identifies entries containing at least one RNA chain in direct contact with a small molecule. The dataset includes structures determined by X-ray crystallography, NMR spectroscopy, and cryo-electron microscopy. A structure was retained if at least one small molecule made direct contact with an RNA nucleotide, defined as any non-hydrogen atom located within 4.8 Å of an RNA heavy atom. Entries were excluded if the ligand simultaneously contacted protein residues, to eliminate complexes in which binding geometry is protein-mediated rather than RNA-intrinsic, or if the ligand appeared on a predefined exclusion list comprising metal ions (e.g., Mg²⁺, K⁺, and Zn²⁺), crystallization additives (e.g., polyethylene glycol fragments, sulfate, and phosphate), and non-pharmacological cofactors (e.g., amino acids or nucleotides) present as crystallization aids rather than as RNA-binding ligands (Supplementary Table S1). For structures containing multiple biological assemblies, only the first assembly was used to avoid coordinate duplication; for NMR ensembles, only the first conformer was submitted for DSSR annotation. This procedure yielded 1,098 PDB entries containing 1,137 RNA–ligand binding events.

Each structure was assigned to one of six functional categories (G-quadruplexes, regulatory RNA motifs, ribosomal RNA, riboswitches, ribozymes, and synthetic aptamers) based on evolutionary origin, functional mechanism, and the structural character of the binding pocket according to predefined keywords (Supplementary Table S2). Category assignment followed a priority hierarchy: structures containing a G-quadruplex binding pocket were assigned to the G-quadruplex category regardless of their biological origin or broader functional context, as G-quadruplex topology defines a structurally and mechanistically distinct recognition mode that is orthogonal to the other five categories. Among the remaining structures, assignments followed the hierarchy: ribosomal RNA, riboswitches, ribozymes, synthetic aptamers, and regulatory RNA motifs. Common RNA motifs not classified under the other five categories form the default “regulatory RNA motifs” category for structured RNAs of diverse biological origin. Category assignments were independently cross-checked against Rfam family classifications (17).

### DSSR-enabled interaction parameter extraction

For each RNA–ligand entry, DSSR v2.9.0 was applied to the atomic coordinate file to generate a JSON-formatted annotation record. DSSR extracted interaction parameters through four sequential stages: (i) annotation of all RNA secondary and tertiary structural elements in the input coordinate file; (ii) identification of RNA nucleotides within 4.8 Å of any ligand heavy atom; (iii) quantification of per-nucleotide interaction types; and (iv) annotation of the tertiary structural context of each contacted nucleotide. The diverse structural parameters were organized into four classes as follows.

Hydrogen-bond topology parameters included the total count of hydrogen bonds between the ligand and RNA (hbonds), resolved by RNA moiety contacted, including the nucleobase (with_base), ribose 2′-OH (with_sugar), and phosphate (with_po4), and classified by donor–acceptor atom-pair (atom_pairs) type (N:N, N:O, and O:O). Partitioning hydrogen bonds by chemical moiety is critical because 2′-OH contacts encode ribose geometry and distinguish RNA from DNA targets, yet are conflated with base contacts in standard contact-counting approaches.

Stacking geometry parameters included the count of RNA nucleotides stacked against the ligand (num_stacked_nts) and the buried stacking surface area (dSASA_stacked_nts, Å²), calculated as the change in solvent-accessible surface area (ΔSASA) upon ligand binding. Here, the complex is restricted to the ligand and its contacting RNA nucleotides. The calculation includes only the ligand atoms, RNA base atoms, and C1′ atoms, explicitly excluding all other RNA backbone atoms. The identity and atom-level detail of each stacked nucleotide were additionally recorded (stacked_nts). This approach distinguishes deep flat intercalation from tangential surface contact that produces comparable nucleotide counts.

Binding pocket descriptors included the total count of contacted RNA nucleotides (contacts), the total buried contact surface area (dSASA_contacted_residues, Å²), an embedding classification (embedded) assigned when the ligand is stacked by RNA bases on both sides, and a groove-preference classification (major_groove_binder or minor_groove_binder).

A ligand was classified as a major-groove binder under three conditions, based on its contacting heavy atoms: (i) all contacting atoms lie on the major groove side; (ii) the number of atoms on the major groove side exceeds those on the minor groove side by ≥ 6; and (iii) the ratio of the number of atoms on the major versus minor groove sides exceeds 3.0. Conversely, a ligand was classified as a minor-groove binder if it met the corresponding reciprocal criteria.

### Unsupervised clustering

The DSSR-derived interaction parameters were compiled into a feature matrix of 1,137 ligand-binding events. Ligand molecular weight and RNA chain length were excluded from the matrix to prevent size confounding. Features were normalized using StandardScaler (z-score standardization: mean centering with unit-variance scaling) to place all interaction parameters on a common scale (18). Principal component analysis (PCA) was then applied to reduce dimensionality; the first 15 principal components, retaining 98.1% of the total variance, were carried forward for K-means clustering with n_init = 50 and random_state = 42 to ensure reproducibility. The optimal cluster number k = 6 was selected from within the elbow region (k = 5–8) based on elbow analysis of the within-cluster sum of squares (Supplementary Figure S1). For two-dimensional visualization, t-distributed stochastic neighbor embedding (t-SNE) was applied to the PCA-reduced matrix with perplexity = 45 and max_iter = 1,000. The statistical association between cluster identity and RNA functional category was assessed by a chi-square test. Pairwise comparisons across functional categories used Kruskal–Wallis omnibus testing, followed by two-sided Mann–Whitney U tests with Benjamini–Hochberg false discovery rate correction (Supplementary Table S3), which were performed in Python using scipy and statsmodels packages (18–20).

### Composite Binding Quality Score

The Composite Binding Quality Score (CBQS) is a seven-metric composite score designed to quantify RNA–ligand interaction quality independently of RNA functional category and ligand size. Seven metrics were selected to represent three mechanistically orthogonal contact-type pairs: hydrogen-bond quality and density (M1, M2), stacking quality and density (M3, M4), and backbone contact quality and density (M5, M6), together with a pocket structural complexity term (M7). Each metric M1–M7 is computed from raw interaction counts, then min-max normalized across all 1,137 SPIRAL ligand-binding events to the [0, 1] range, where 0 corresponds to the minimum and 1 to the maximum value observed in the dataset:

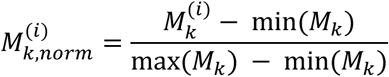

where *k* is the metric number, and *i* is the entry number. The CBQS for entry i is then the unweighted arithmetic mean of all seven normalized sub-scores, multiplied by 100 to express the result on a 0–100 scale:

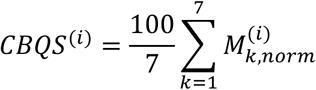

The individual metrics are defined as follows: M1 = num_hbonds / num_contacts (hydrogen-bond quality), where num_hbonds is the count of RNA–ligand hydrogen bonds classified as standard or acceptable quality by DSSR’s donor–acceptor distance and angle criteria; M2 = num_hbonds / (dSASA_contacted_residues / 100) (hydrogen-bond density per 100 Å^2^ buried area); M3 = num_stacked_nts / num_contacts (stacking quality); M4 = dSASA_stacked_nts / dSASA_contacted_residues (stacking density); M5 = (with_sugar + with_po4) / num_contacts (backbone contact quality); M6 = (with_sugar + with_po4) / (dSASA_contacted_residues / 100) (backbone contact density per 100 Å² buried area); and M7 = sum of per-contact counts across seven tertiary motif features (hairpin loop, junction loop, pseudoknot, splayed-apart, coaxial stack, multiplet, and non-canonical pair; pocket structural complexity). A CBQS of 100 represents the theoretical maximum in which all seven sub-scores simultaneously achieve the highest value observed in the dataset. Category-level CBQS values reported in Table 1 are the means of entry-level scores within each functional category.

**Table 1.** CBQS scores and sub-score profiles across six RNA functional categories.

| Category | CBQS | M1 | M2 | M3 | M4 | M5 | M6 | M7 |
| --- | --- | --- | --- | --- | --- | --- | --- | --- |
| Riboswitches | 32.2 | 34.0 | 34.2 | 23.6 | 47.9 | 21.0 | 23.5 | 41.1 |
| Ribozymes | 28.9 | 32.3 | 32.6 | 18.3 | 37.5 | 20.2 | 22.7 | 38.4 |
| Synthetic aptamers | 26.1 | 21.7 | 17.4 | 38.4 | 53.3 | 13.4 | 11.7 | 26.8 |
| G-quadruplexes | 24.1 | 8.7 | 6.8 | 43.5 | 58.1 | 8.8 | 7.0 | 35.6 |
| Ribosome | 23.0 | 31.3 | 23.5 | 12.8 | 19.4 | 21.4 | 18.9 | 33.4 |
| Regulatory RNA motifs | 19.4 | 15.1 | 9.8 | 35.6 | 42.5 | 12.5 | 8.6 | 11.8 |

### Binding affinity dataset and analysis

Experimental binding affinity data were obtained from PDBbind (version NL.2020R1) (21) and supplemented by manual literature curation for RNA–ligand structures deposited after 2020, yielding 275 entries with affinity measurements. All values were converted to a unified *pK_d_* scale:

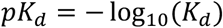

where *K_d_* is expressed in molar units, such that a higher *pK_d_* corresponds to tighter binding. *K_i_* and *IC₅₀* values were treated as approximate *K_d_* equivalents following established practice (22).

Affinity–feature correlation analyses were performed on the 275 SPIRAL entries with a curated, standardized *pK_d_* value. Spearman rank correlation was used to assess monotonic associations between individual interaction parameters and *pK_d_* (Supplementary Table S4). A two-predictor linear regression model incorporating total buried contact surface area (dSASA_contacted_residues) and C2′-endo pucker count as RobustScaler-normalized features was fitted to this dataset and evaluated by five-fold cross-validation (20 repeats), with predictive performance reported as the mean cross-validated R² (R² = 0.40). All statistical analyses were performed in Python using the scipy (v1.17.1), scikit-learn (v1.8.0), and statsmodels (v0.14) packages (18–20), with all p-values being two-sided and a significance threshold of α = 0.05 applied throughout.

## RESULTS

### SPIRAL dataset composition

Application of the curation criteria described in the Methods to the full PDB yielded 1,098 entries containing 1,137 RNA–ligand binding events, determined by X-ray crystallography, NMR spectroscopy, and cryo-electron microscopy (Figure 1A). The dataset spans PDB structures deposited through May 2026, with depositions accelerating over the past decade, reflecting the cryo-EM resolution revolution for large ribosomal complexes (23) alongside the growing structural characterization of riboswitches and regulatory RNA motifs (Figure 1B). For each entry, DSSR-derived interaction parameters are provided alongside functional category assignments, cluster identities, and CBQS scores.

**Figure 1.**
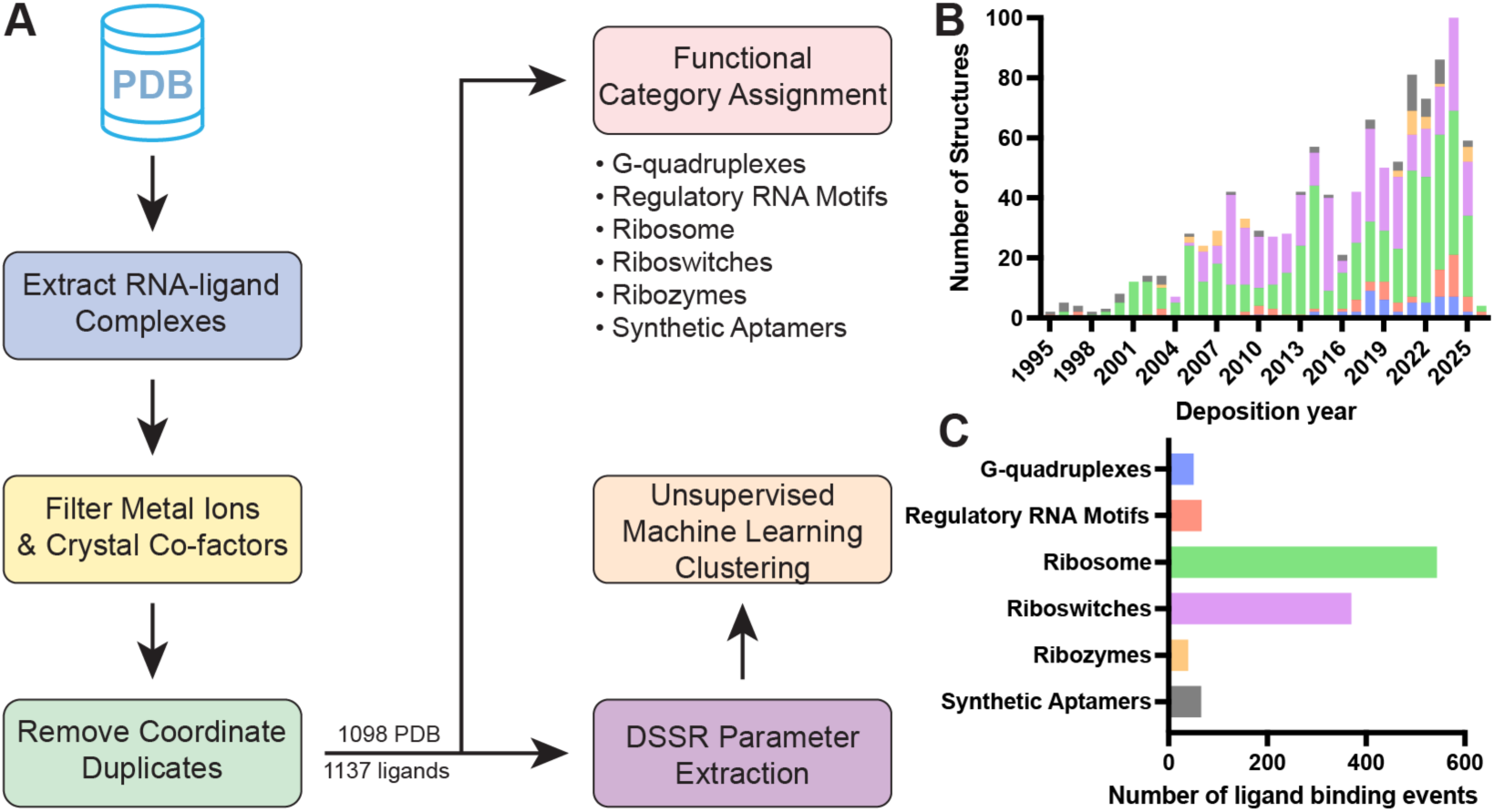
SPIRAL dataset curation workflow and composition. A) Curation pipeline: PDB-wide search for RNA–small molecule structures, followed by sequential exclusion of metal ions, buffer components, and protein-contacting ligands, and the retention of the first biological assembly, yielding 1,098 entries and 1,137 binding events. B) Growth of SPIRAL entries over time by RNA functional category. C) Distribution of ligand-binding events across six RNA functional categories; colors match those used in panel B. The six category colors assigned here, G-quadruplexes (blue), regulatory RNA motifs (coral), ribosomal RNA (green), riboswitches (lavender), ribozymes (gold), and synthetic aptamers (grey), are used consistently in all subsequent figures.

The 1,137 binding events were distributed across six functional RNA categories defined by evolutionary origin, functional mechanism, and the structural character of the binding pocket (Figure 1C). Ribosomal RNA represents the largest category (n = 543, 47.8%), comprising ribosomal RNA domains and subdomains targeted primarily by antibiotics, followed by riboswitches (n = 370, 32.5%), which are natural metabolite sensors that undergo ligand-induced conformational changes to regulate gene expression. Synthetic aptamers (n = 66, 5.8%) are SELEX-derived or engineered RNA constructs selected for high-affinity ligand recognition *in vitro*. RNA G-quadruplexes (n = 51, 4.5%) are four-stranded G-rich structures whose stacked G-tetrad surfaces are recognized by planar aromatic compounds. Ribozymes (n = 40, 3.5%) are catalytic RNAs whose active sites bind substrate analogues, cofactors, or inhibitors. The remaining 67 structures (5.9%) comprise regulatory RNA motifs, a heterogeneous category encompassing disease-relevant structural elements, including viral regulatory elements, trinucleotide repeat expansions, and stem-loop structures, that do not fall within the other five categories. Together, these six categories span the major architectural classes of functional RNA.

### A DSSR-enabled pipeline for RNA–small molecule interaction profiling

Existing structural analysis tools designed for protein–ligand systems lack the capacity to describe the tertiary RNA features that constitute small-molecule binding interfaces. To address this limitation, we developed the “ligand” module for DSSR v2.9.0, extending the core annotation engine with new interaction-analysis capabilities built on the standard base reference frame (24).

Because this frame is rigidly defined relative to each Watson-Crick pair, with its x-axis pointing toward the major groove, a ligand’s position can be expressed in intrinsic structural coordinates rather than absolute space, providing a principled geometric basis for classifying the binding mode.

The groove-preference feature assigns each complex a groove-binder classification from the distribution of ligand contacts across the major and minor groove edges. In the base-pair reference frame, atoms contacting the major-groove side carry positive x-coordinates and those contacting the minor-groove side carry negative x-coordinates, so the groove preference follows directly from the signed coordinates of the ligand’s contacting atoms. Figure 2A illustrates this for a major-groove binder bound to the HIV-1 frameshift-site RNA (PDB ID: 2L94), in which the contacting ligand atoms carry large positive x-coordinates, unambiguously placing the ligand on the major-groove face. This captures a binding-mode distinction not accessible from contact counts alone. Risdiplam bound to the SMN2 5′-splice site illustrates this classification for a therapeutically relevant ligand: despite forming only two hydrogen bonds, both are directed to the major-groove face of A14 (25), classifying it as a major-groove binder (Supplementary Figure S2).

**Figure 2.**
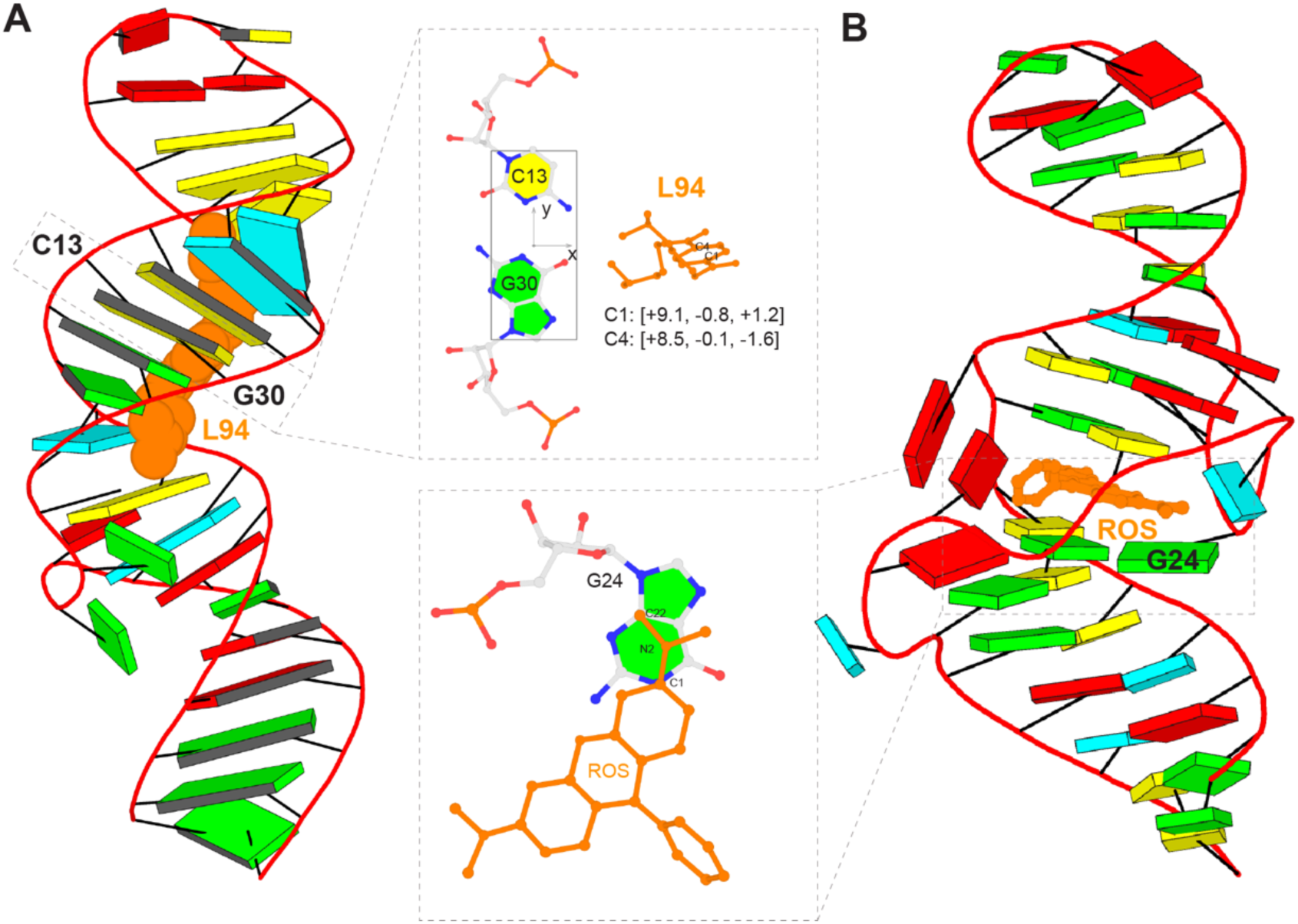
Algorithms in the DSSR “ligand” module for identifying groove-binding and stacking-binding ligands. A) Groove detection, illustrated with a major-groove binder bound to the HIV-1 frameshift-site RNA (PDB ID: 2L94). The left panel shows the full structure with base pairs as schematic blocks (with minor-groove edges in black) and the ligand L94 in an orange space-filling representation. The top-middle inset panel shows the standard base-pair reference frame on the C13–G30 base pair. B) Stacking detection, illustrated with the malachite green aptamer bound to tetramethyl-rosamine (ROS; PDB ID: 1F1T). The right panel shows the full structure with bases as schematic blocks and the embedded ligand in orange stick representation; the bottom-middle inset panel shows ROS stacking on G24. In panels A and B, the RNAs are depicted as red ribbons, with bases and Watson–Crick base pairs represented as color-coded blocks: A and A-U in red, C and C-G in yellow, G and G-C in green, and U and U-A in cyan.

Given the exceptional diversity in ligand geometry, molecular size, and planarity, the stacking feature quantifies RNA–ligand stacking using the rigidly defined nucleobase as a coordinate reference. A stacking contact is identified when any ligand heavy atom lies within 3.6 Å of a base ring system, geometrically extended to include exocyclic groups and the ribose C1′ atom (Figure 2B). Because a single ligand can contact one base through several atoms and can engage multiple bases across a pocket, the module records stacking at atomic resolution and computes the buried stacking surface area by differential solvent-accessible surface area, providing a continuous measure that distinguishes deep flat intercalation from tangential surface contact producing comparable nucleotide counts.

The embedding feature classifies whether a ligand is enclosed by RNA on both sides through stacking. When coordinating bases sandwich the ligand from geometrically opposing faces, it is classified as embedded, satisfying the spatial criteria for a classic intercalator binding mode, as illustrated by tetramethyl-rosamine in the malachite green aptamer (PDB ID 1F1T; Figure 2B). Furthermore, adenosylcobalamin bound to its cognate riboswitch (26) exemplifies another embedded class, with its corrin ring enclosed by 22 contacted nucleotides and penetrating to the groove floor rather than docking at the surface (Supplementary Figure S2).

The hydrogen-bond feature additionally detects and classifies all standard RNA–ligand hydrogen bonds by the RNA moiety contacted (nucleobase, ribose 2′-OH, or phosphate), enabling 2′-OH recognition to be tracked independently of base-specific contacts. Together, these capabilities operate on the same atomic coordinate framework as DSSR’s existing tertiary structure annotation, ensuring that each interaction parameter is reported in its full tertiary structural context and generating the interaction fingerprint that is the analytical foundation of SPIRAL.

### Unsupervised clustering resolves six mechanistically distinct binding modes partitioned by RNA functional class

K-means clustering of the SPIRAL entries resolved six mechanistically distinct binding modes. The t-SNE projection revealed clear spatial separation among the clusters (Figure 3A), and the association between cluster identity and RNA functional category was highly significant (χ² test, χ² = 1428, p < 0.001), confirming that RNA functional class is a strong determinant of binding mode identity (Figure 3B).

**Figure 3.**
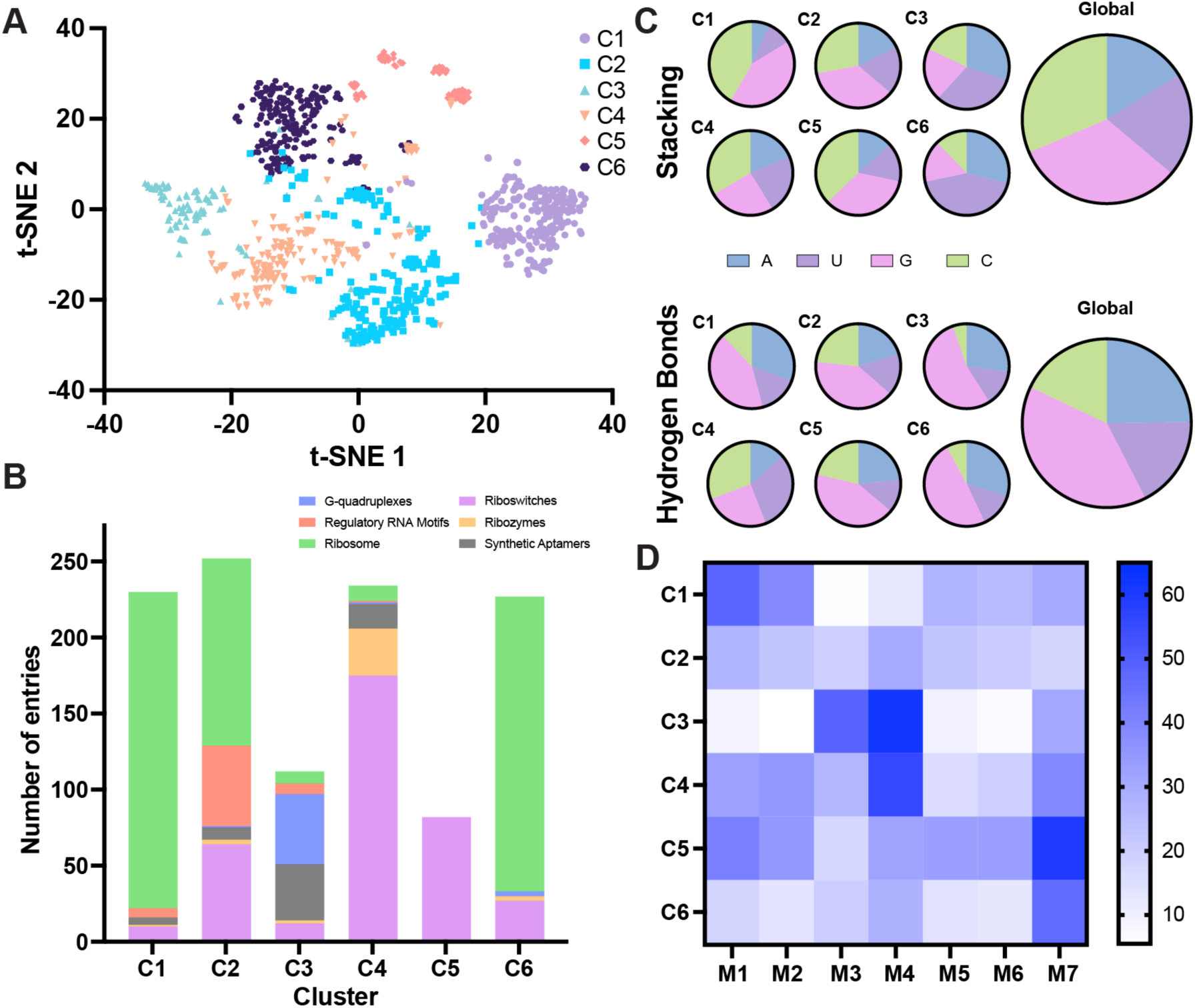
Unsupervised clustering of SPIRAL RNA–ligand complexes into six mechanistically distinct binding modes. A) t-SNE projection of the PCA-reduced feature matrix (15 components, 98.1% variance retained), colored by cluster identity (C1–C6). B) RNA functional category composition of each cluster; the strong association between cluster identity and RNA functional class was confirmed by chi-square test (p < 0.001). The six category colors assigned here, G-quadruplexes (blue), regulatory RNA motifs (coral), ribosomal RNA (green), riboswitches (lavender), ribozymes (gold), and synthetic aptamers (grey), are used consistently in all subsequent figures. C) Per-nucleotide frequency of hydrogen-bond and stacking contacts summed across all ligand-binding events in the SPIRAL dataset. D) Heatmap of cluster centroid values for seven interaction parameters (M1–M7); color intensity reflects the normalized score on a 0–65 scale.

Each of the six clusters is defined by dominant interaction features. Cluster 1 (C1) is an electrostatic ribosomal cluster (90% ribosomal RNA) with the highest hydrogen-bond quality and density and the highest phosphate-backbone and coaxial-stack contacts of any cluster, but near-zero RNA–ligand stacking, consistent with the open-groove aminoglycoside A-site recognition mechanism. C2 is a mixed, moderate-engagement cluster of intermediate character, populated by ribosomal RNA (49%), riboswitches (25%), and regulatory RNA motifs (21%). C3 is a stacking-dominated cluster co-populated by G-quadruplex (41%) and synthetic aptamer (33%) binders, which are united by the highest stacking quality and density and the lowest hydrogen-bond quality of any cluster, reflecting aromatic intercalation and G-tetrad stacking modes; it also shows the highest C2′-endo enrichment, consistent with the distinctive sugar puckers of these folds. C4 is a riboswitch-dominant cluster (75% riboswitch, 13% ribozyme) with the highest pseudoknot and hairpin-loop contacts and strong stacking density, consistent with engagement of multi-helix architectures across diverse tertiary folds. C5 is an exclusively riboswitch cluster (100%) with the highest backbone contact quality and density, pocket complexity, coaxial-stack contacts, sugar-2′-OH contacts, and non-canonical base-pair contacts, reflecting engagement of the deepest and most structurally complex riboswitch pockets. C6 is dominated by ribosomal RNA (85%), with uniformly few interactions across the hydrogen-bond and backbone dimensions despite the highest junction-loop contacts, reflecting predominantly surface-docking interactions without deep tertiary fold penetration. Across all six clusters, guanosine was the most frequently contacted nucleotide for both hydrogen-bond and stacking interactions (Figure 3C), consistent with the dual chemical character of guanine as both a hydrogen-bond donor/acceptor and the largest aromatic stacking surface among the four canonical nucleobases (22,27).

Statistical analysis confirmed globally significant stratification of interaction parameters across the six functional categories (Supplementary Table S3), with post-hoc pairwise comparisons identifying riboswitches and regulatory RNA motifs as the most consistently separated pair across individual metrics. Three category-specific profiles are particularly informative for RNA-targeted drug design.

Riboswitches exhibited the highest hydrogen-bond quality and density, the highest 2′-OH contact fraction, and the highest pocket structural complexity of all six categories (Table 1), collectively reflecting the multi-point recognition strategy by which riboswitches discriminate between structurally similar metabolites at sub-μM affinity (28,29). Among all RNA functional classes, minor-groove contacts constitute a primary recognition strategy only in riboswitches, where base-edge hydrogen bonds and 2′-OH interactions simultaneously read nucleobase identity and ribose geometry, providing higher information content per contact than stacking or phosphate interactions alone.

Synthetic aptamers displayed a systematic stacking bias relative to riboswitches, with stacking quality and density significantly higher and hydrogen-bond density significantly lower. This pattern likely reflects the structural preference of SELEX for intercalation-competent RNA architectures: flat, aromatic small molecules readily intercalate into solvent-accessible internal loops and three-way junctions, a mode known to dominate selection campaigns and requiring active counter-selection to suppress (30,31).

Ribosomal RNA binders had the highest phosphate-contact fraction of any category, consistent with the electrostatic complementarity mechanism of aminoglycosides, and were distributed across five clusters with structurally distinct interaction parameter profiles (Figure 3B). Drug class distribution across the two largest ribosome sub-sites further validates this structural segregation (Supplementary Figure S3). The ribosomal RNA binders within Cluster 1 (R-C1, n = 208) are dominated by aminoglycosides at the 30S decoding A-site (98% of its binders), characterized by the highest hydrogen-bond quality, the highest phosphate and coaxial-stack contacts, and near-zero RNA–ligand stacking of any cluster (Figure 4A). Cluster 6’s ribosomal RNA binders (R-C6, n = 194) comprise ligands of the 50S peptidyl-transferase center and exit tunnel, including macrolides/ketolides (26%), lincosamides (14%), and other natural-product translation inhibitors, and exhibits the highest junction-loop contacts of any ribosome cluster together with uniformly low hydrogen-bond and backbone scores, consistent with predominantly surface-docking engagement of these secondary sites. The remaining ribosome entries distribute across three additional clusters (R-C2, R-C3, and R-C4) without distinctive interaction signatures, likely reflecting the structural diversity of secondary ribosome binding sites and mixed drug classes such as tetracyclines (Supplementary Figure S3).

**Figure 4.**
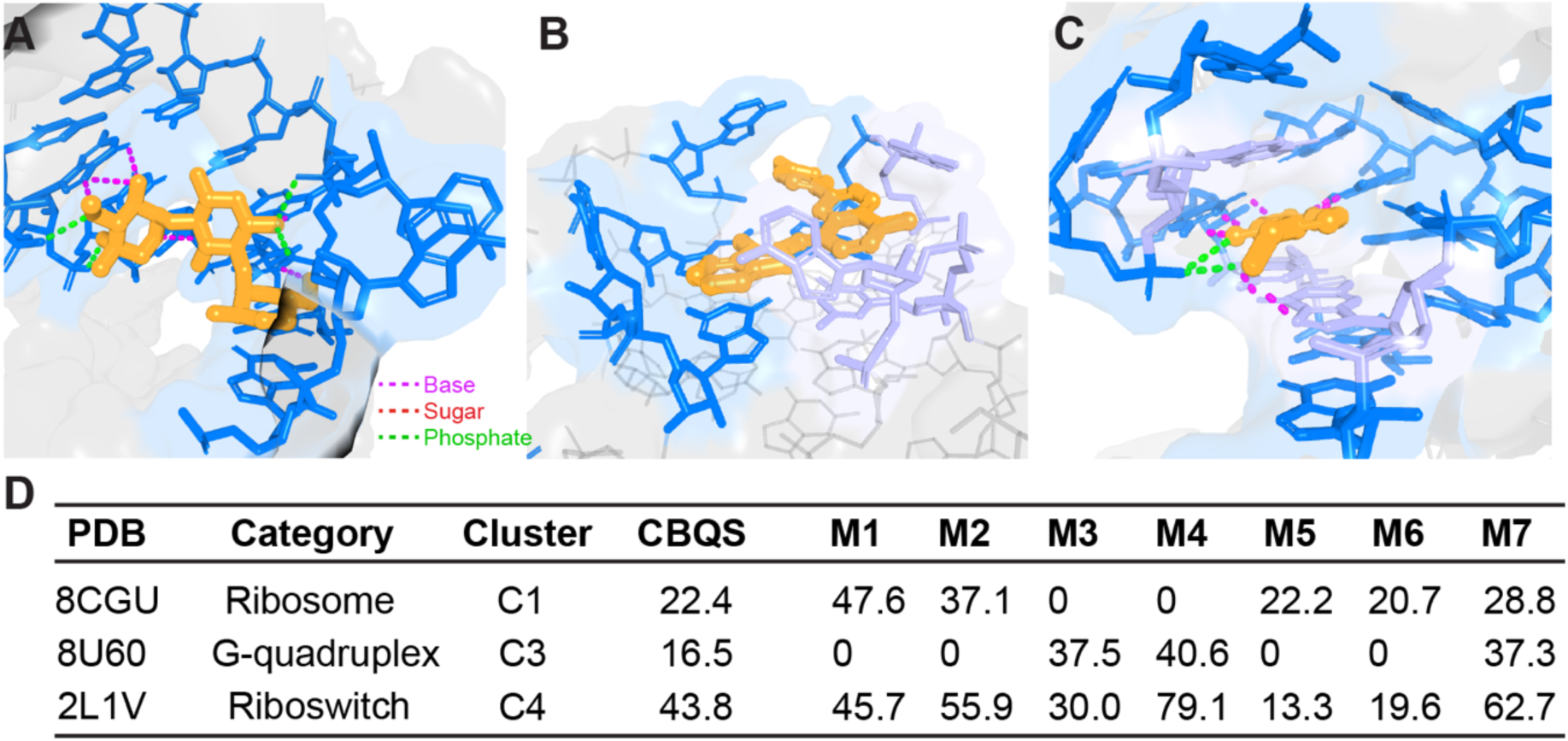
Representative RNA–small molecule binding modes and their CBQS sub-score profiles. A) Ribosomal RNA A-site binding by gentamicin (PDB ID: 8CGU). The ligand engages the decoding site through eight hydrogen-bond contacts to RNA bases and four phosphate contacts, with no stacking interaction. B) RNA G-quadruplex end-stacking by a thiazole orange ligand (PDB ID: 8U60). The ligand sits flat on the external G-tetrad surface through pure aromatic stacking against three nucleotides, with no hydrogen bonds to RNA. C) preQ1 riboswitch minor-groove recognition by preQ1 (PDB ID: 2L1V). The ligand is buried within the pseudoknot pocket and forms ten hydrogen-bond contacts to RNA bases and two phosphate contacts, alongside stacking interactions with three nucleotides. D) CBQS and sub-score profiles for all three structures. In all structural panels: full RNA, faded grey with a transparent grey surface; interface nucleotides, marine; stacking nucleotides, light blue; ligand, orange; hydrogen bonds to RNA bases, magenta dashes; hydrogen bonds to phosphate, green dashes; and hydrogen bonds to sugar, red dashes.

That this structural segregation is recoverable from DSSR interaction parameters alone, without prior knowledge of binding site identity or drug class, demonstrates the discriminatory power of the SPIRAL feature set. G-quadruplex binders were distinguished from all other categories by a near-complete absence of groove preference and a heavy reliance on G-tetrad stacking (Figure 4B).

### CBQS: a seven-metric framework for RNA–ligand binding quality

The six binding mode clusters span fundamentally different chemical strategies, from the dense hydrogen-bond networks of riboswitch minor-groove recognition to the purely geometric enclosure of G-tetrad end-stacking. Single-axis metrics borrowed from protein–ligand analysis, such as hydrogen-bond count or stacking area, cannot distinguish whether equivalent affinity arises from polar recognition, geometric enclosure, or aromatic stacking, and therefore produce quality rankings biased by category composition rather than by the quality of the underlying recognition event. We therefore developed the CBQS, which captures interaction quality across three orthogonal contact-type pairs: hydrogen-bond quality and density (M1/M2), stacking quality and density (M3/M4), and backbone contact quality and density (M5/M6), together with pocket structural complexity (M7), all normalized to the global SPIRAL distribution (see Methods). The distinct centroid profiles across the six clusters (Figure 3D) confirm that M1–M7 capture mechanistically orthogonal dimensions of RNA–ligand recognition: riboswitch-enriched clusters show high hydrogen-bond metrics (M1 and M2) with intermediate stacking; G-quadruplex and SELEX-enriched clusters show the inverse pattern (high M3 and M4; near-zero M1 and M2); and ribosomal clusters are distinguished primarily by hydrogen-bond quality and backbone contact metrics (Figure 5A,B). This variation confirms that M1–M7 together span the full chemical diversity of RNA–ligand recognition in the SPIRAL dataset, providing the empirical justification for their selection as CBQS components. CBQS scores and sub-score profiles for all six functional categories are summarized in Table 1.

**Figure 5.**
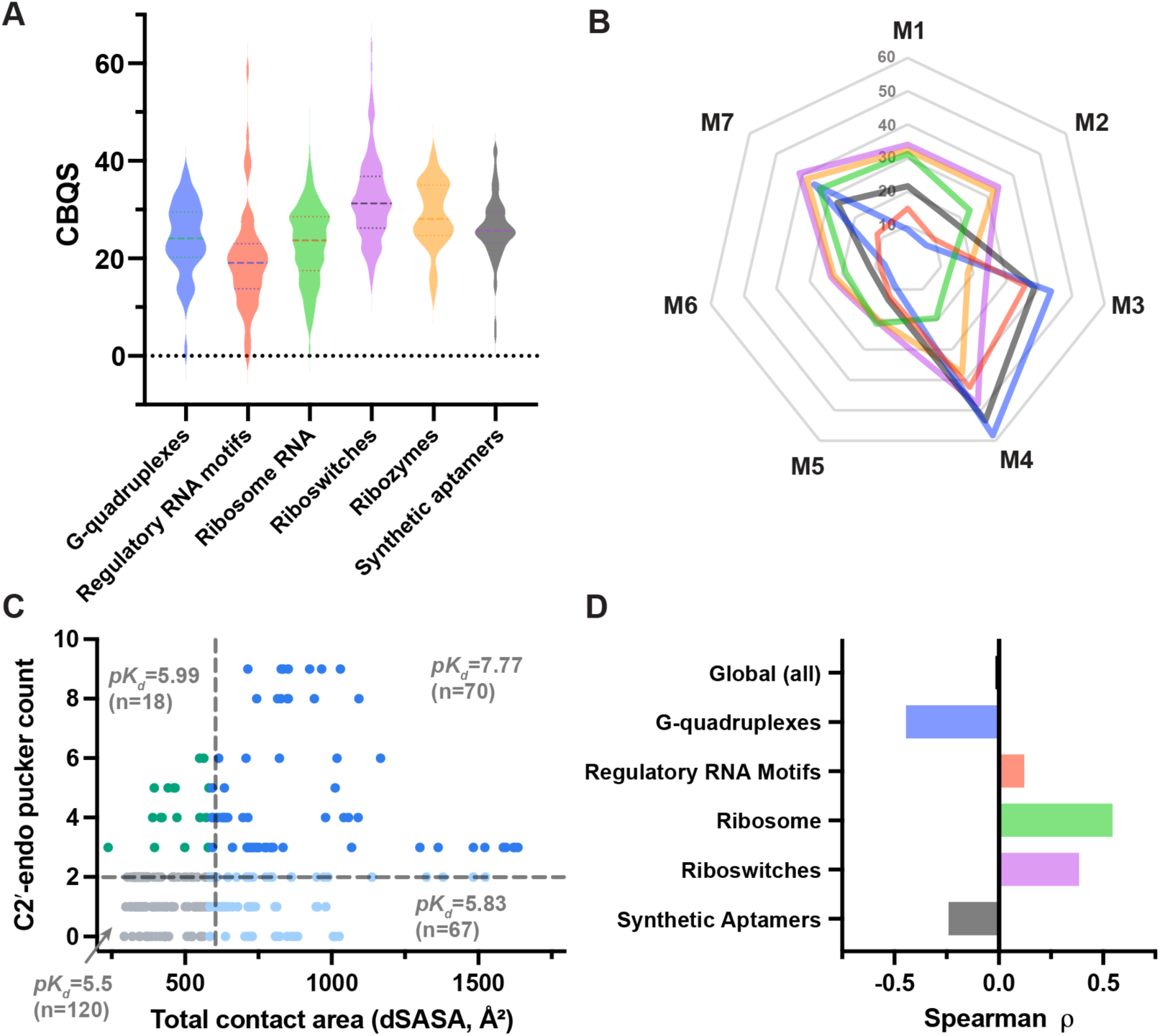
CBQS scores and binding affinity analysis. A) Distribution of CBQS across the six RNA functional categories. Each violin shows the full distribution of per-entry CBQS values within a category. Internal lines mark the median and interquartile range. B) The radar plots show sub-score profiles (M1–M7) for all six functional categories, illustrating mechanistically distinct recognition strategies including the two categories (synthetic aptamers and G-quadruplexes) that achieve statistically equivalent overall CBQS scores. C) Quadrant analysis of mean *pK_d_* across four bins defined by a median split on total buried contact area (median = 583.0 Å²) and C2′-endo pucker count (median = 2.0) for the 275 deduplicated affinity entries; the mean *pK_d_* for each quadrant is shown. D) Category-stratified Spearman rank correlations between hydrogen-bond count and *pK_d_*, illustrating a Simpson’s paradox: the global correlation is non-significant (ρ = −0.016, n = 275), whereas the within-riboswitch correlation is strongly positive (ρ = +0.384, p < 0.001) and the within-G-quadruplex correlation is significantly negative (ρ = −0.444, p = 0.014), reflecting mechanistically opposing roles of hydrogen bonding in polar versus stacking-driven recognition. Ribozymes were excluded from the stratified analysis (n = 1 affinity entry, insufficient for correlation).

Riboswitches ranked highest (CBQS = 32.2), consistent with the multi-point hydrogen-bond recognition strategy that natural aptamers have evolved to achieve metabolite discrimination at sub-μM affinity. Regulatory RNA motifs ranked lowest (CBQS = 19.4), reflecting their predominant surface-docking binding mode and near-complete absence of junction-loop contacts (Table 1). Ribozymes occupied the second rank (CBQS = 28.9), with balanced contributions across all seven metrics.

Synthetic aptamers (CBQS = 26.1) and RNA G-quadruplexes (CBQS = 24.1) occupied statistically indistinguishable positions, yet their sub-score profiles were mechanistically distinct: synthetic aptamers were dominated by the stacking pair (M3/M4) with low hydrogen-bond scores, while G-quadruplex binders showed the highest stacking density of any category (M4 = 58.1) with virtually no hydrogen-bond signal (M1 = 8.7 and M2 = 6.8), consistent with recognition through geometric enclosure at the G-tetrad surface (Figure 5A,B). This result reveals a key limitation of single-number scoring: equal scores can arise from mechanistically incompatible recognition strategies, and the sub-score breakdown is required to distinguish them. These archetypes, together with representative structural examples, are illustrated in Figure 4. Ribosomal RNA ranked fifth (CBQS = 23.0), consistent with the open-groove, phosphate-electrostatic binding mode that dominates this category.

Despite having the highest CBQS scores, riboswitches were distributed across all six binding mode clusters, reflecting genuine mechanistic diversity among their aptamer families. The largest riboswitch sub-population resides in Cluster 4 (C4, n = 175, 47%), the junction/complex-pocket cluster, followed by Cluster 5 (C5, n = 82, 22%), the exclusively riboswitch cluster representing the deepest and most complex binding pockets in the dataset, and Cluster 2 (C2, n = 64, 17%), the mixed moderate-engagement cluster (Figure 3B). The remaining riboswitches are distributed across C1, C3, and C6.

Riboswitches are the only RNA functional class in SPIRAL in which minor-groove contacts constitute a primary recognition strategy. Minor-groove recognition is chemically distinctive: contacts to the groove floor simultaneously read nucleobase identity through base-edge hydrogen bonds and ribose geometry through 2′-OH interactions, providing higher information content per contact than either stacking or phosphate interactions alone. It is this property that accounts for the elevated hydrogen-bond quality and density scores (M1 = 34.0, M2 = 34.2) that distinguish riboswitches from all other categories in the CBQS sub-score analysis.

### Regulatory RNA motifs binders exhibit the lowest CBQS score with five quantifiable design gaps

Regulatory RNA motifs ranked lowest in CBQS among all six functional categories. Unlike riboswitches, whose ligands are natural metabolites shaped by co-evolution with the aptamer fold, or ribosomal RNA binders, which are predominantly antibiotics optimized through decades of medicinal chemistry, the majority of ligands in the regulatory RNA motifs category were identified through *in vitro* binding assays or cell-based functional screens and have not undergone equivalent structural optimization. Structurally, regulatory RNA motif binders are characterized by high coaxial-stack contacts and moderate stacking density, but substantially lower hydrogen-bond density, 2′-OH contact fraction, and pocket complexity than riboswitches and ribozymes.

Five specific design gaps were identified by comparing the regulatory RNA motif interaction parameter profile against the riboswitch benchmark (Table 2). Junction-loop contacts were absent from every regulatory RNA motif entry, representing the most severe gap. Pseudoknot contacts, hairpin-loop contacts, base-multiplet contacts, and ribose 2′-OH contacts were each substantially underrepresented relative to riboswitches, with fold-gaps ranging from 6.6–57.8. Regulatory RNA motif binders show the highest coaxial-stack contacts of any category (mean 3.81 per entry), indicating that ligands in this class already engage helix–helix interfaces but do not extend into the flanking loop architectures where the highest-information-content contacts occur. No current regulatory RNA motif entry in SPIRAL satisfies all five gap criteria simultaneously, providing a specific benchmark for future ligand design campaigns targeting this RNA class. Whether engaging these underrepresented structural sites would improve binding affinity is addressed in the following section.

**Table 2.** Five quantifiable design gaps for regulatory RNA motif binders relative to the riboswitch benchmark.

| Design gap | Regulatory motifs | Riboswitch | Fold gap |
| --- | --- | --- | --- |
| Junction loop | 0 | 4.346 | ∞ |
| Pseudoknot | 0.015 | 0.862 | 57.8X |
| Hairpin-loop | 0.194 | 2.941 | 15.2X |
| Multiplet | 0.597 | 4.992 | 8.4X |
| 2'-OH | 0.358 | 2.370 | 6.6X |

These are mean values per functional category

### C2′-endo pucker count and total buried contact area predict binding affinity

To identify the structural interaction parameters that govern RNA–ligand binding affinity, Spearman rank correlation was applied to the 275-entry affinity dataset. C2′-endo sugar pucker count and total buried contact surface area emerged as the two strongest predictors of *pK_d_* (ρ = +0.530 and +0.440, respectively, both p < 0.001; Supplementary Table S4), together explaining 40% of *pK_d_* variance in a two-predictor linear regression model (Figure 5C). The two predictors are only weakly intercorrelated (ρ = +0.31), indicating that they capture largely complementary structural contributions to affinity.

The mechanistic basis of the C2′-endo sugar pucker count association is instructive. C3′-endo is the thermodynamically preferred ribose conformation in A-form RNA; switching to C2′-endo widens the local groove, reorients the nucleobase, and repositions the 2′-OH axially, at an energetic cost that is only recoverable when RNA–ligand complementarity is sufficiently strong. C2′-endo sugar pucker count therefore serves as a proxy for induced-fit binding depth rather than a simple geometric descriptor of the pocket. C2′-endo enrichment in the SPIRAL dataset was concentrated at junction-loops, non-canonical base-pair, and multiplet-network bases. This is consistent with fundamental structural studies showing that mixed C2’-endo / C3’-endo sugar puckering is a strict biophysical requirement for stabilizing highly conserved, non-canonical architectures like the GpU dinucleotide platform and the loop-E motif, where the rigid conformation swap establishes specialized backbone-mediated interaction edges (32). This convergence between the strongest affinity predictor and the design gaps implicates a specific class of RNA pocket, one characterized by tertiary structural strain, loop-based geometry, and high C2′-endo enrichment, as the highest-priority structural target class for regulatory RNA motif optimization.

Hydrogen-bond count showed no significant global correlation with *pK_d_* (Figure 5D), a result that is resolved as a Simpson’s paradox upon category-stratified analysis (Supplementary Table S4). Within riboswitches, which contribute 61% of the affinity dataset (n = 169), hydrogen-bond count was strongly and positively correlated with *pK_d_* (ρ = +0.384; Supplementary Table S4), consistent with the multi-point polar recognition strategy of riboswitch aptamers. Within G-quadruplexes, the same correlation was negative and statistically significant (ρ = −0.444), reflecting the geometric, stacking-driven recognition mode of these binders, in which added polar substituents impose an entropic penalty without commensurate binding gain.

Pooling categories with opposing within-category relationships produced the observed near-zero global correlation. This finding has a direct practical consequence: hydrogen-bond optimization strategies validated within one RNA functional class cannot be assumed to transfer to another, and affinity models must be applied within category-specific structural frameworks.

Total stacking buried area correlated positively with *pK_d_* globally (ρ = +0.458), whereas the stacking fraction did not (ρ = +0.075; non-significant). This dissociation indicates that it is the absolute size of the buried contact footprint, rather than its chemical composition, that governs affinity. Increasing aromatic bulk therefore improves potency through a non-specific contact-area effect while simultaneously reducing selectivity by displacing the base-specific hydrogen-bond contacts that encode sequence specificity, a trade-off consistent with the observed tendency of high-stacking RNA binders to accumulate off-target liabilities in cellular contexts (4,33).

## DISCUSSION

### SPIRAL provides tertiary structural context for RNA–small molecule recognition

Several databases of RNA–small molecule interactions have been established, each addressing a distinct aspect of RNA–ligand recognition. R-BIND 2.0 (11,12) provides comprehensive chemical annotation of bioactive RNA-targeting ligands and links them to RNA secondary structure motifs, serving as a cheminformatics reference for ligand property analysis. HARIBOSS (13) compiles RNA–ligand structures from the PDB with pocket geometry descriptors derived from three-dimensional coordinates, providing a structural complement to R-BIND. Inforna (14) enables sequence-based matching of RNA structural motifs to known binding small molecules, supporting rational ligand identification from RNA sequence information. DRLiPS (34) predicts druggable RNA pockets from structural data using machine learning, extending pocket analysis to unliganded RNA targets. SPIRAL complements these resources by providing, to our knowledge, the first database in which every RNA–ligand entry carries DSSR-computed tertiary structural context annotation across four classes of interaction parameters, enabling systematic binding-mode classification and category-independent binding quality scoring that are not addressed by existing resources.

Prior structural analyses have found that RNA pockets are, on average, less hydrophobic and more solvent-exposed than protein binding sites (13,22). The SPIRAL dataset qualifies this generalization: the picture is not uniform across RNA functional classes. Riboswitch pockets achieved embedding rates of 72% and hydrogen-bond densities approaching those reported for protein–ligand interfaces, whereas regulatory RNA motif binding sites were highly solvent-exposed and poorly enclosed. The CBQS score difference between these two categories (32.2 vs. 19.4) quantifies this structural heterogeneity. It argues against treating RNA as a single target class in screening library design or computational pocket detection, where category-specific structural criteria are needed.

### Structural determinants of RNA–ligand affinity and selectivity

C2′-endo pucker count is the strongest global affinity predictor across 275 structurally diverse complexes. To our knowledge, this is the first demonstration of a quantitative affinity correlation for this conformational feature in the small-molecule–RNA field. C2′-endo nucleotides are highly overrepresented in catalytic active sites and critical tertiary structures, exhibit slow local conformational dynamics with half-lives on the 10–100 s timescale in native tertiary context, and have been proposed to function as molecular timers in RNA folding and ligand recognition (35). The energetic cost of the C3′-endo to C2′-endo transition is only offset when ligand–RNA complementarity is sufficient to recover it, making C2′-endo sugar pucker count a structural indicator of induced-fit binding depth rather than a geometric descriptor. Independent support comes from protein–RNA structural studies showing that C2′-endo enrichment is significantly concentrated at interfaces bound by sequence-specific protein domains and that C2′-endo nucleotides are preferentially involved in direct recognition contacts (36), suggesting that C2′-endo enrichment is a general property of high-affinity RNA recognition sites rather than a phenomenon specific to any one class of binding partner.

The convergence of the affinity and CBQS design gap analyses on the same structural sites, including junction loops, non-canonical base pairs, and multiplet networks, is the central mechanistic conclusion of this study. These elements impose geometric strain that favors C2′-endo ribose conformations, and the multi-point contacts they present are inherently sequence-specific. Designing regulatory RNA motif binders that engage these sites is therefore predicted to improve both potency and selectivity concurrently. The dissociation between total stacking buried area and stacking fraction reinforces this: absolute contact footprint governs affinity, not chemical composition. Adding aromatic bulk to increase stacking area improves binding affinity through a size-dependent contact effect that is indifferent to RNA sequence and therefore gains potency at the cost of selectivity, a trade-off documented across multiple RNA target classes (4,33).

The mechanistic divergence between polar and stacking recognition strategies, quantified by the CBQS sub-score profiles, is further confirmed by the opposing within-category hydrogen-bond correlations (Table 1). The stacking dominance of SELEX-derived aptamers likely reflects the enrichment of intercalation-competent architectures during in vitro selection rather than an intrinsic ceiling on recognition quality, since natural evolution converged on minor-groove contacts that simultaneously read nucleobase identity and 2′-OH geometry, providing the chemical specificity needed to discriminate between metabolites at sub-μM affinity. Synthetic aptamers achieve a high overall CBQS score approaching that of riboswitches (26.1 vs. 32.2) largely through elevated stacking scores (M3 = 38.4 and M4 = 53.3), while their hydrogen-bond quality score (M1 = 21.7) is roughly two-thirds that of riboswitches (M1 = 34.0), implying that screening strategies biased toward minor-groove contacts will yield leads with improved selectivity (Table 1).

### Design principles and applications for RNA-targeted drug discovery

The five design gaps in regulatory RNA motif binders (Table 2) translate into specific structural criteria. The complete absence of junction-loop contacts in every regulatory RNA motif entry is the most severe gap, yet this category ranks first in coaxial-stack contacts, indicating that current ligands already engage helix–helix interfaces. One approach is to extend a functional group from the coaxial-stack anchor into the flanking junction or hairpin loop, the two largest loop-architecture gaps. The gaps in pseudoknot and multiplet contacts indicate that penetration to the tertiary fold depth where the highest-specificity contacts occur requires three-dimensional, non-planar ligand architectures. No current regulatory RNA motif entry satisfies all five criteria simultaneously, providing a specific structural benchmark for future optimization campaigns.

Beyond synthetic design, the CBQS sub-scores provide a mechanistic complement to docking scores for virtual screening hit prioritization. Poses that recapitulate the contact-type balance of the target RNA cluster can be prioritized over mechanistically inconsistent ones. For a riboswitch target, this means favoring high 2′-OH contacts and junction-loop engagement; for a G-quadruplex, high stacking density. In structure-guided optimization, the sub-scores diagnose which contact types are underrepresented relative to the category benchmark: a hit with high stacking density but low hydrogen-bond quality and near-zero 2′-OH contacts should be elaborated with functional groups capable of minor-groove hydrogen bonding rather than additional aromatic bulk.

### Summary

SPIRAL provides a systematic meta-analysis of RNA–small molecule recognition built on DSSR-computed tertiary interaction parameters. Three principal findings emerge. First, unsupervised clustering resolves six mechanistically distinct binding modes whose distribution is strongly governed by RNA functional class, demonstrating that tertiary structural context determines small-molecule engagement in ways that secondary structure alone cannot capture. Second, the CBQS framework reveals that equivalent binding quality is achieved through mechanistically distinct strategies, and the gap between the highest- and lowest-scoring categories reflects quantifiable structural deficits that translate into actionable design targets. Third, the C2′-endo pucker count and total buried contact area are the two dominant affinity predictors; their enrichment at the same tertiary structural sites most under-engaged by current regulatory RNA motif binders establishes a structurally convergent criterion for optimizing potency and selectivity simultaneously. CBQS sub-scores further provide a mechanistic complement to docking scores for virtual screening hit prioritization and a diagnostic tool for structure-guided optimization, both of which fall outside the scope of standard scoring functions calibrated on protein–ligand systems.

## ACKNOWLEDGEMENTS

The research reported in this publication was supported by the National Institute of General Medical Sciences (NIGMS) of the National Institutes of Health under Award Number R24GM153869 to X.J.L. and Medical College of Wisconsin Research Affairs Committee New Faculty Pilot Grant to Y.W. We thank the members of the Wang lab for helpful discussions.

## AUTHOR CONTRIBUTIONS

Yaqiang Wang: Conceptualization, Formal analysis, Methodology, Visualization, and Writing. Xiang-Jun Lu: Conceptualization, Formal analysis, Methodology, and Writing.

## DATA AVAILABILITY

The SPIRAL database is freely available at https://www.wangrnalab.org/spiral, providing an interactive structure viewer together with all curated entries and their DSSR-derived interaction parameters, RNA functional category assignments, cluster assignments, and CBQS scores.

Structural interaction parameters were computed with the ligand module of DSSR v2.9.0, which is freely available to academic users at https://x3dna.org. All structural coordinates analyzed are available from the Protein Data Bank (https://www.rcsb.org) under their respective accession codes. Experimental binding affinities were obtained from PDBbind (NL.2020R1) and from the primary literature.

## REFERENCES

1. Childs-Disney, J.L., Yang, X., Gibaut, Q.M.R., Tong, Y., Batey, R.T. and Disney, M.D. (2022) Targeting RNA structures with small molecules. Nature reviews. Drug discovery, 21, 736–762.

2. Disney, M.D., Dwyer, B.G. and Childs-Disney, J.L. (2018) Drugging the RNA World. Cold Spring Harbor perspectives in biology, 10.

3. Henley, M.J. and Koehler, A.N. (2021) Advances in targeting ‘undruggable’ transcription factors with small molecules. Nature reviews. Drug discovery, 20, 669–688.

4. Warner, K.D., Hajdin, C.E. and Weeks, K.M. (2018) Principles for targeting RNA with drug-like small molecules. Nature reviews. Drug discovery, 17, 547–558.

5. Haniff, H.S., Knerr, L., Liu, X., Crynen, G., Bostrom, J., Abegg, D., Adibekian, A., Lekah, E., Wang, K.W., Cameron, M.D. et al. (2020) Design of a small molecule that stimulates vascular endothelial growth factor A enabled by screening RNA fold-small molecule interactions. Nature chemistry, 12, 952–961.

6. Rzuczek, S.G., Southern, M.R. and Disney, M.D. (2015) Studying a Drug-like, RNA-Focused Small Molecule Library Identifies Compounds That Inhibit RNA Toxicity in Myotonic Dystrophy. ACS chemical biology, 10, 2706–2715.

7. Baranello, G., Darras, B.T., Day, J.W., Deconinck, N., Klein, A., Masson, R., Mercuri, E., Rose, K., El-Khairi, M., Gerber, M. et al. (2021) Risdiplam in Type 1 Spinal Muscular Atrophy. N. Engl. J. Med., 384, 915–923.

8. Ratni, H., Ebeling, M., Baird, J., Bendels, S., Bylund, J., Chen, K.S., Denk, N., Feng, Z., Green, L., Guerard, M. et al. (2018) Discovery of Risdiplam, a Selective Survival of Motor Neuron-2 (SMN2) Gene Splicing Modifier for the Treatment of Spinal Muscular Atrophy (SMA). J. Med. Chem., 61, 6501–6517.

9. Veenbaas, S.D., Felder, S. and Weeks, K.M. (2026) fpocketR: A Platform for Identification and Analysis of Ligand-Binding Pockets in RNA. ACS chemical biology, 21, 151–159.

10. Veenbaas, S.D., Koehn, J.T., Irving, P.S., Lama, N.N. and Weeks, K.M. (2025) Ligand-binding pockets in RNA and where to find them. Proc Natl Acad Sci U S A, 122, e2422346122.

11. Donlic, A., Swanson, E.G., Chiu, L.Y., Wicks, S.L., Juru, A.U., Cai, Z., Kassam, K., Laudeman, C., Sanaba, B.G., Sugarman, A. et al. (2022) R-BIND 2.0: An Updated Database of Bioactive RNA-Targeting Small Molecules and Associated RNA Secondary Structures. ACS chemical biology, 17, 1556–1566.

12. Morgan, B.S., Sanaba, B.G., Donlic, A., Karloff, D.B., Forte, J.E., Zhang, Y. and Hargrove, A.E. (2019) R-BIND: An Interactive Database for Exploring and Developing RNA-Targeted Chemical Probes. ACS chemical biology, 14, 2691–2700.

13. Panei, F.P., Torchet, R., Menager, H., Gkeka, P. and Bonomi, M. (2022) HARIBOSS: a curated database of RNA-small molecules structures to aid rational drug design. Bioinformatics, 38, 4185–4193.

14. Disney, M.D., Winkelsas, A.M., Velagapudi, S.P., Southern, M., Fallahi, M. and Childs-Disney, J.L. (2016) Inforna 2.0: A Platform for the Sequence-Based Design of Small Molecules Targeting Structured RNAs. ACS chemical biology, 11, 1720–1728.

15. Lu, X.J., Bussemaker, H.J. and Olson, W.K. (2015) DSSR: an integrated software tool for dissecting the spatial structure of RNA. Nucleic Acids Res., 43, e142.

16. Li, S., Olson, W.K. and Lu, X.J. (2019) Web 3DNA 2.0 for the analysis, visualization, and modeling of 3D nucleic acid structures. Nucleic Acids Res., 47, W26–W34.

17. Ontiveros-Palacios, N., Cooke, E., Nawrocki, E.P., Triebel, S., Marz, M., Rivas, E., Griffiths-Jones, S., Petrov, A.I., Bateman, A. and Sweeney, B. (2025) Rfam 15: RNA families database in 2025. Nucleic Acids Res., 53, D258–D267.

18. Pedregosa, F., Varoquaux, G., Gramfort, A., Michel, V., Thirion, B., Grisel, O., Blondel, M., Prettenhofer, P., Weiss, R., Dubourg, V. et al. (2011) Scikit-learn: Machine Learning in Python. J Mach Learn Res, 12, 2825–2830.

19. Virtanen, P., Gommers, R., Oliphant, T.E., Haberland, M., Reddy, T., Cournapeau, D., Burovski, E., Peterson, P., Weckesser, W., Bright, J. et al. (2020) SciPy 1.0: fundamental algorithms for scientific computing in Python. Nat Methods, 17, 261–272.

20. Seabold, S. and Perktold, J. (2010), Proceedings of the 9th Python in Science Conference.

21. Liu, Z., Li, Y., Han, L., Li, J., Liu, J., Zhao, Z., Nie, W., Liu, Y. and Wang, R. (2015) PDB-wide collection of binding data: current status of the PDBbind database. Bioinformatics, 31, 405–412.

22. Padroni, G., Patwardhan, N.N., Schapira, M. and Hargrove, A.E. (2020) Systematic analysis of the interactions driving small molecule-RNA recognition. RSC Med Chem, 11, 802–813.

23. Kuhlbrandt, W. (2014) Biochemistry. The resolution revolution. Science, 343, 1443–1444.

24. Olson, W.K., Bansal, M., Burley, S.K., Dickerson, R.E., Gerstein, M., Harvey, S.C., Heinemann, U., Lu, X.J., Neidle, S., Shakked, Z. et al. (2001) A standard reference frame for the description of nucleic acid base-pair geometry. J. Mol. Biol., 313, 229–237.

25. Malard, F., Wolter, A.C., Marquevielle, J., Morvan, E., Ecoutin, A., Rudisser, S.H., Allain, F.H.T. and Campagne, S. (2024) The diversity of splicing modifiers acting on A-1 bulged 5’-splice sites reveals rules for rational drug design. Nucleic Acids Res., 52, 4124–4136.

26. Johnson, J.E., Jr., Reyes, F.E., Polaski, J.T. and Batey, R.T. (2012) B12 cofactors directly stabilize an mRNA regulatory switch. Nature, 492, 133–137.

27. Conner, A.V., Kim, L.M., Fagan, P.A., Harding, D.P. and Wheeler, S.E. (2025) Stacking Interactions of Druglike Heterocycles with Nucleobases. J Chem Inf Model, 65, 3502–3516.

28. Kavita, K. and Breaker, R.R. (2023) Discovering riboswitches: the past and the future. Trends Biochem. Sci., 48, 119–141.

29. Breaker, R.R. (2022) The Biochemical Landscape of Riboswitch Ligands. Biochemistry, 61, 137–149.

30. Ellington, A.D. and Szostak, J.W. (1990) In vitro selection of RNA molecules that bind specific ligands. Nature, 346, 818–822.

31. Ruscito, A. and DeRosa, M.C. (2016) Small-Molecule Binding Aptamers: Selection Strategies, Characterization, and Applications. Front Chem, 4, 14.

32. Lu, X.J., Olson, W.K. and Bussemaker, H.J. (2010) The RNA backbone plays a crucial role in mediating the intrinsic stability of the GpU dinucleotide platform and the GpUpA/GpA miniduplex. Nucleic Acids Res., 38, 4868–4876.

33. Costales, M.G., Childs-Disney, J.L., Haniff, H.S. and Disney, M.D. (2020) How We Think about Targeting RNA with Small Molecules. J. Med. Chem., 63, 8880–8900.

34. Krishnan, S.R., Roy, A., Wong, L. and Gromiha, M.M. (2025) DRLiPS: a novel method for prediction of druggable RNA-small molecule binding pockets using machine learning. Nucleic Acids Res., 53.

35. Mortimer, S.A. and Weeks, K.M. (2009) C2’-endo nucleotides as molecular timers suggested by the folding of an RNA domain. Proc Natl Acad Sci U S A, 106, 15622–15627.

36. Kligun, E. and Mandel-Gutfreund, Y. (2015) The role of RNA conformation in RNA-protein recognition. RNA biology, 12, 720–727.

