## Supplemental file for "Structural Pockets and Interacting RNA-Associated Ligands (SPIRAL): A DSSR-enabled Meta-Analysis of RNA-Small Molecule Recognition"

**Figure S1**

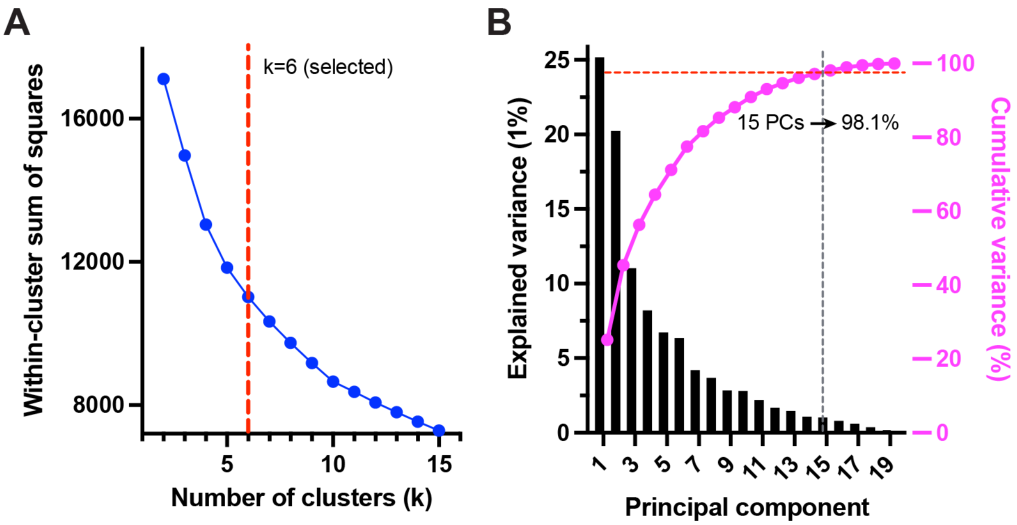


**Figure S1 Elbow analysis for k selection.** A) Elbow analysis. Within-cluster sum of squares plotted for k = 2 to 15, with k = 6 marked by a dashed red line. B) PCA explained variance: individual component variance (19 components) shown as bars with cumulative variance as a red overlay line, with the 98.1% threshold marked, confirming that 15 components capture 98.1% of variance with negligible information loss.

**Figure S2**

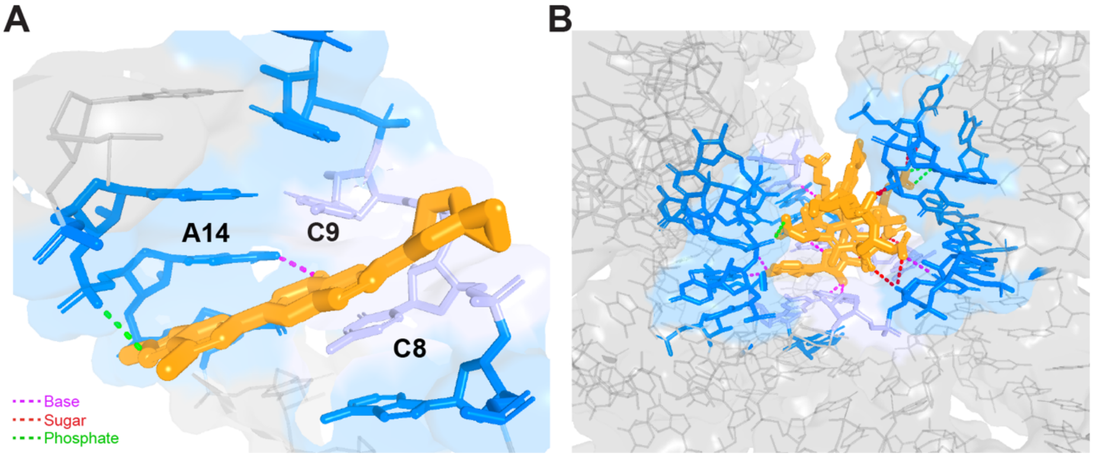


**Figure S2. Representative RNA–small molecule structures illustrating DSSR-computed interaction parameters extracted by SPIRAL.** A) Risdiplam (PDB ID: 8R62) bound to the SMN2 pre-mRNA 5′-splice site duplex. Its binding mode exemplifies the major groove recognition class: the ligand interacts with two cytosine residues (C8 and C9) within the RNA stem. Only two hydrogen bonds are formed: one to the base of A14 (N6) and one to its phosphate (OP2), while no 2′-OH contacts are detected, and the binding site is classified as a major groove binder. B) Adenosylcobalamin (PDB ID: 4GMA) bound to the adenosylcobalamin riboswitch. The large corrin ring system is enclosed by 22 contacted nucleotides spanning junction loops, hairpin loops, a pseudoknot, and kissing-loop elements, with a total buried contact surface area of 1867 Å$^2$. Five nucleotides stack on the corrin ring, and 17 hydrogen bonds are distributed across all three RNA moiety classes: six to nucleobases, five to phosphate oxygens, and six to ribose 2′-OH groups. In both panels, the full RNA is shown as faded grey sticks with a transparent grey surface; interface nucleotides are marine; stacking nucleotides are light blue; the ligand is orange; hydrogen bonds to bases, phosphate oxygens, and ribose 2′-OH groups are shown as magenta, green, and red dashes, respectively.

**Figure S3**

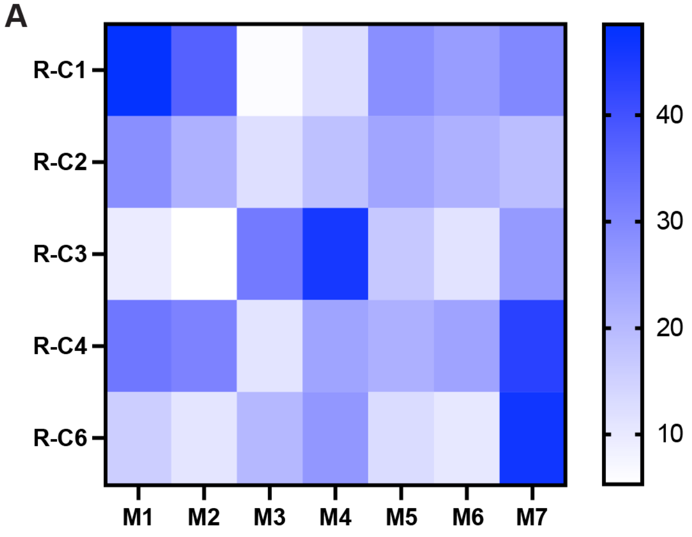


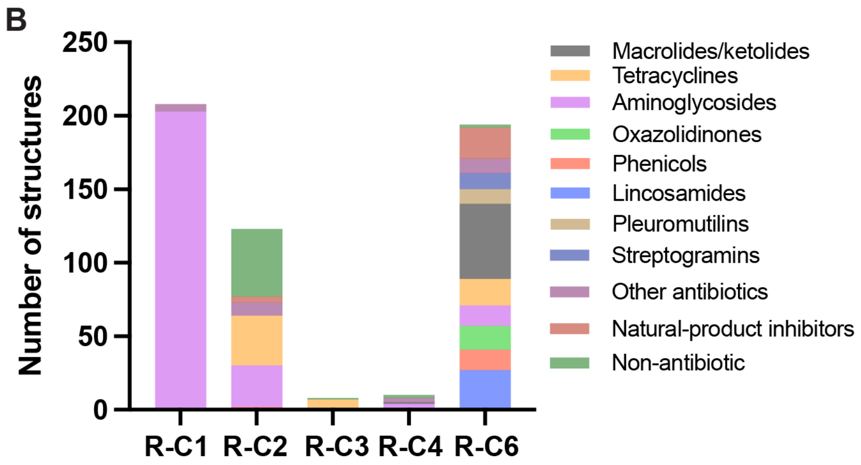


**Figure S3 Interaction profiles and drug class distribution across five ribosome RNA binding mode sub-sites.** A) Heatmap of interaction parameter profiles for the five clusters containing ribosome RNA entries (R-C1, R-C2, R-C3, R-C4, and R-C6; C5 is excluded as it contains no ribosome RNA entries). Rows represent sub-sites sorted by entry count; columns represent seven CBQS sub-scores (M1–M7); color intensity reflects the normalized score on a 0–50 scale. B) Approximate drug class distribution across the five ribosome sub-sites, expressed as percentage of ligand-binding events per cluster. R-C1 is dominated by aminoglycosides (98%), corresponding to the 30S decoding A-site. R-C6 groups 50S peptidyl-transferase-center and exit-tunnel antibiotics, including macrolides and ketolides (26%), lincosamides (14%), and oxazolidinones (8%). R-C2 is a mixed cluster combining non-antibiotic ligands (37%), tetracyclines (28%), and aminoglycosides (23%). R-C3 is enriched for tetracyclines (75%).

**Table S1 Summary of the predefined exclusion list applied during SPIRAL dataset curation**

| Functional category | Amount | CCD codes |
|---|---|---|
| Metal ions and elemental atoms | 28 | BA, BR, C, CA, CD, CL, CO, 3CO, CS, CU1, F, HG, IRI, K, LU, MG, MN, N, NA, NH4, O, OS, PB, RHD, SR, TL, W, ZN |
| Inorganic ions and clusters | 14 | FES, FS2, NCO, NO3, OHX, PO4, POP, SE4, SF4, SO4, VTR, VTE, WO2, WO4 |
| PEG fragments | 11 | 1PE, 2PE, 4PE, 5PE, 6PE, 7PE, 8PE, P6G, PEG, PG4, PGE |
| Buffer components | 14 | ACE, ACT, ACY, 3TD, BTB, CAC, EPE, GZ6, MES, MLI, N2P, N3D, TLA, TRS |
| Cryoprotectants and solvents | 11 | BME, DMS, EDO, EOH, GOL, HEZ, IPA, IPH, MPD, PDI, TER |
| Other crystallization additives | 15 | 2HP, B3P, CPT, DPO, GAI, GDE, IHP, NME, PUT, S9L, SPD, SPK, SPM, TAM, URE |
| Non-pharmacological cofactors | 3 | ARF, FME, SIN |
| Nucleotide-related compounds | 10 | 8OS, 50L, 50N, A1CGJ, EQ1, EQ4, G3A, GP3, LXI, LXR |
| Amino acids | 7 | ARG, GLY, LYS, MET, PHE, PRO, TYR |
| Other non-drug-like biomolecules | 5 | 6MZ, CAD, FRU, GLC, SEY |
| Unidentified PDB placeholders | 2 | UNL, UNX |
| Total | 120 | |

**Table S2 Functional RNA PDB category assignment criteria for the SPIRAL dataset**

| Category | Amount* | % | Keywords | Rfam family |
|---|---|---|---|---|
| G-quadruplexes | 50 | 4.6% | quadruplex, G4, G-tetrad, G-quartet, fluorescent, dimer, biotin, hexammine, iridium, Spinach, Mango, Corn, Beetroot, Chili | Not Rfam-classified (structural class) |
| Regulatory motifs | 67 | 6.1% | TAR, bulge, HIV, interaction, hairpin, stem-loop, solution NMR | RF00250 (HIV-1 TAR); and other functional RNA motif families |
| Ribosome | 512 | 46.6% | ribosomal, subunit, rRNA, ribosome, translation, mitochondrial, cytosolic | RF00177 (small subunit rRNA); RF02543 (large subunit rRNA); RF00001 (5S rRNA) |
| Riboswitches | 368 | 33.5% | riboswitch, aptamer, domain, variant, mutant, ligand-gated | RF00050 (FMN); RF00059 (TPP); RF00162 (SAM); RF00167 (purine); RF00174 (cobalamin); RF01739 (glutamine); RF01786 (c-di-GMP); RF00522 (preQ1); RF01734 (fluoride) |
| Ribozymes | 38 | 3.5% | ribozyme, snRNP, splicing, substrate, spliceosome, self-splicing, ribonucleoprotein, glmS | RF00163 (hammerhead); RF00173 (hairpin); RF01577 (RNase P); RF02001 (group II intron) |
| Synthetic aptamers | 63 | 5.7% | aptamer, Fab, chain, heavy, light, theophylline, soak, SELEX, Pepper, Squash, RhoBAST | Not Rfam-classified (in vitro selected constructs) |
| Total | 1098 | 100% | | |

*Amount = PDB entries (structures), whereas n values in main text are ligand binding events.

### Table S3 Statistical analyses of cluster and category comparisons

| Comparison | Test | Statistic | n | p / q value | Sidedness | Significant (α=0.05) |
|---|---|---|---|---|---|---|
| Cluster identity × RNA category | Chi-square test of independence | $\chi^2 = 1428.4$, df = 25 | 1137 | 3.32e-286 | two-sided | Yes |
| Overall CBQS across 6 categories | Kruskal–Wallis omnibus | H = 258.4 | 1137 | 8.60e-54 | two-sided | Yes |
| H-bond quality (M1) across 6 categories | Kruskal–Wallis omnibus | H = 146.4 | 1137 | 7.68e-30 | two-sided | Yes |
| Stacking density (M4) across 6 categories | Kruskal–Wallis omnibus | H = 422.0 | 1137 | 5.45e-89 | two-sided | Yes |
| Pocket complexity (M7) across 6 categories | Kruskal–Wallis omnibus | H = 218.6 | 1137 | 3.03e-45 | two-sided | Yes |
| Riboswitches vs Ribosome (CBQS) | Mann–Whitney U (BH-FDR) | U = 156915 | 370 / 543 | q < 0.001 | two-sided | Yes |
| Riboswitches vs Regulatory motifs (CBQS) | Mann–Whitney U (BH-FDR) | U = 21306 | 370 / 67 | q < 0.001 | two-sided | Yes |
| Riboswitches vs G-quadruplexes (CBQS) | Mann–Whitney U (BH-FDR) | U = 14223 | 370 / 51 | q < 0.001 | two-sided | Yes |
| Riboswitches vs Synthetic aptamers (CBQS) | Mann–Whitney U (BH-FDR) | U = 17352 | 370 / 66 | q < 0.001 | two-sided | Yes |
| Ribozymes vs Regulatory motifs (CBQS) | Mann–Whitney U (BH-FDR) | U = 2188 | 40 / 67 | q < 0.001 | two-sided | Yes |
| Synthetic aptamers vs Regulatory motifs (CBQS) | Mann–Whitney U (BH-FDR) | U = 3433 | 66 / 67 | q < 0.001 | two-sided | Yes |
| Ribozymes vs Ribosome (CBQS) | Mann–Whitney U (BH-FDR) | U = 15404 | 40 / 543 | q < 0.001 | two-sided | Yes |
| Ribosome vs Regulatory motifs (CBQS) | Mann–Whitney U (BH-FDR) | U = 23654 | 543 / 67 | q < 0.001 | two-sided | Yes |
| G-quadruplexes vs Regulatory motifs (CBQS) | Mann–Whitney U (BH-FDR) | U = 2373 | 51 / 67 | q < 0.001 | two-sided | Yes |
| Ribozymes vs G-quadruplexes (CBQS) | Mann–Whitney U (BH-FDR) | U = 1395 | 40 / 51 | q = 0.004 | two-sided | Yes |

| Comparison | Test | Statistic | n | p / q value | Sidedness | Significant (α=0.05) |
|---|---|---|---|---|---|---|
| Synthetic aptamers vs Ribosome (CBQS) | Mann–Whitney U (BH-FDR) | U = 21950 | 66 / 543 | q = 0.004 | two-sided | Yes |
| Ribozymes vs Synthetic aptamers (CBQS) | Mann–Whitney U (BH-FDR) | U = 1650 | 40 / 66 | q = 0.040 | two-sided | Yes |
| Riboswitches vs Ribozymes (CBQS) | Mann–Whitney U (BH-FDR) | U = 8743 | 370 / 40 | q = 0.068 | two-sided | No |
| Synthetic aptamers vs G-quadruplexes (CBQS) | Mann–Whitney U (BH-FDR) | U = 1950 | 66 / 51 | q = 0.153 | two-sided | No |
| G-quadruplexes vs Ribosome (CBQS) | Mann–Whitney U (BH-FDR) | U = 15074 | 51 / 543 | q = 0.295 | two-sided | No |

Symbols: $\chi^2$, chi-square statistic; df, degrees of freedom; H, Kruskal–Wallis statistic; U, Mann–Whitney U statistic; q, Benjamini–Hochberg FDR-adjusted p value; α, significance threshold. All tests are two-sided. For pairwise comparisons, n is given as the size of each group (group A / group B). Chi-square evaluated cluster identity against RNA functional category across 1,137 binding events; Kruskal–Wallis tests compared each interaction metric across the six categories; pairwise category comparisons used Mann–Whitney U tests with Benjamini–Hochberg correction. Analyses were performed in Python using scipy and statsmodels.

**Table S4 Affinity correlation statistics (n = 275 affinity-characterized entries)**

| Analysis | Variable / subgroup | Spearman ρ | 95% CI | n | p value | Significant (α=0.05) |
|---|---|---|---|---|---|---|
| Global predictors | C2′-endo pucker count | +0.530 | [+0.44, +0.61] | 275 | 2.49e-21 | Yes |
| Global predictors | Buried contact area (Å²) | +0.440 | [+0.34, +0.53] | 275 | 1.76e-14 | Yes |
| Global predictors | H-bond count | -0.016 | [-0.13, +0.10] | 275 | 7.88e-01 | No |
| Simpson's paradox (H-bond vs pKd) | Global (pooled) | -0.016 | [-0.13, +0.10] | 275 | 0.788 | No |
| Simpson's paradox (H-bond vs pKd) | Riboswitches | +0.384 | [+0.25, +0.51] | 169 | 2.5e-07 | Yes |
| Simpson's paradox (H-bond vs pKd) | Synthetic aptamers | -0.241 | [-0.53, +0.09] | 36 | 0.156 | No |
| Simpson's paradox (H-bond vs pKd) | G-quadruplexes | -0.444 | [-0.69, -0.10] | 30 | 0.014 | Yes |
| Simpson's paradox (H-bond vs pKd) | Ribosome RNA | +0.544 | [+0.04, +0.83] | 15 | 0.036 | Yes |
| Simpson's paradox (H-bond vs pKd) | Regulatory motifs | +0.122 | [-0.30, +0.50] | 24 | 0.571 | No |

Symbols: ρ, Spearman rank correlation coefficient; CI, confidence interval. All correlations are two-sided Spearman rank correlations; 95% confidence intervals were computed by Fisher z-transformation. Ribozymes (n = 1 affinity entry) were not estimable and are omitted. A two-predictor linear regression model (buried contact area and C2′-endo pucker count, RobustScaler-normalized) achieved a mean five-fold cross-validated $R^2$ of 0.40. Analyses were performed in Python using scipy and scikit-learn.
